# Phenotypic and transcriptional features of the Araliaceae species under distinct light environments

**DOI:** 10.1101/2022.08.22.504876

**Authors:** Yu-Qian Niu, Yu-Xin Zhang, Xin-Feng Wang, Zhen-Hui Wang, Ji Yang, Yu-Guo Wang, Wen-Ju Zhang, Zhi-Ping Song, Lin-Feng Li

## Abstract

Elucidating how the plant species respond to variable light conditions is essential to understand the ecological adaptation to heterogeneous environments. Plant performance and gene regulatory network underpinning the adaptation have been well-documented in sun-grown species. In this study, we surveyed phenotypic and transcriptomic features of four shade-grown and one sun-grown woody species of the family Araliaceae under distinct light conditions. Our phenotypic comparisons demonstrate that the four shade-grown species possess lower light saturation point and higher assimilation ability of the net photosynthetic rate compared to the sun-grown species. In particular, the four shade-grown species maintain similar photosynthesis efficiency in both highlight and lowlight conditions. However, a significantly decreased photosynthesis rate was observed under lowlight condition of the sun-grown species compared to highlight condition. In addition, our leaf anatomical analyses reveal that while all the five species show different anatomical structures under distinct light conditions, the shade-grown species possess lower degree of phenotypic plasticity relative the sun-grown species. Further comparisons of the transcriptome profiling show that all the five species exhibit gene expression divergence among different light conditions. The differentially expressed genes identified in the five species are functionally related to photosynthesis, secondary metabolites and other basic metabolisms. More importantly, differential regulation of the photosynthesis- and photomorphogenesis-related genes are potentially correlated with the phenotypic responses to distinct light conditions of the five species. Our study provides new insights on how the sun- and shade-grown woody species respond to shade and sunlight environments.

## INTRODUCTION

Plants are sessile by nature that have evolved developmental and phenotypic plasticity to respond to heterogeneous environments (Sultan, 2000; Sullivan & Deng, 2003; Delagrange et al., 2004; Bou-Torrent et al., 2012). Sunlight is one of the most important environmental factors that not only serves as an energy source for photosynthesis but also acts as signal to regulate plant development and growth (Lau & Deng, 2010). However, the light quantity and quality vary dramatically in nature both on temporal and spatial scales (Hudson et al., 2017). To survive, plant species have developed sophisticated mechanisms to sense dynamic light environments and evoke a suite of short- and long-term responses (Sullivan & Deng, 2003; Bou-Torrent et al., 2012; Huber et al., 2021; Liu et al., 2021). In open habitats, heliophytic (sun-grown) plants receive full light signaling components (*i*.*e*., red, blue and UV-B) to regulate developmental and physiological activities (Fiorucci & Fankhauser, 2017). When shaded, sun-grown plants perceive shade signals (*i*.*e*., low red/far-red ratio) and express shade avoidance syndrome (SAS) to escape the micro-environments, such as elongation of stem and petiole, reducing branches, and accelerated flowering (Casal, 2013; Fiorucci & Fankhauser, 2017; Gommers, 2017).

Sciophytic (shade-grown) plants that grown in the forest understory developed shade-tolerance phenotype to cope with limited light environments (Baltzer & Thomas, 2007; Valladares & Niinemets, 2008). Compared to the SAS that has developed as an escape mechanism, the shade-grown plants adopt a shade-tolerance strategy to complete entire life cycle in shaded habitat (Gommers et al., 2013). Three hypotheses have been proposed on how shade-grown plants adapt to shaded habitat, including carbon gain, stress tolerance and phenotypic plasticity (Valladares & Niinemets, 2008). The carbon gain hypothesis holds that shade-tolerance refers to the maximizing light capture and use in photosynthesis and with minimizing respiratory costs (Givnish, 1988). Under this hypothesis, any morphological and physiological traits that can improve light utilization efficiency and carbon acquisition are beneficial to the environmental adaptability (Givnish, 1988). However, the shade tolerance hypothesis argues that the most critical determinant for a shade-grown species is to improve the ability of resistance to biotic and abiotic stress rather than invest a large proportion of energy to promote plant growth and development (Augspurger, 1984; Kitajima, 1994). Comparisons of the shade-avoidance and defense-response traits confirmed that shade-tolerant plants tend to allocate more energy to storage organs instead of the elongation of stem and petiole under low light condition (Kobe, 1997; Canham, 1999; Ballare & Austin, 2019). In addition, the hypothesis of phenotypic plasticity proposes that shade-tolerant species are expected to show low degree of physiological and morphological plasticity under diverse light conditions (Valladares et al., 2002). Indeed, meta-analyses of more than 70 traits revealed that woody and shade-tolerant species have low level of plasticity at the majority of the analyzed traits, such as photosynthetic capacity and realized growth (Poorter et al., 2019). This empirical evidence suggest that shade-tolerance is not solely the suppression of classic SAS but also a complicated process of ecological adaptation to shade environment. However, it still remained under-investigated of how woody shade-tolerant species respond to variable light conditions.

The family Araliaceae consists of approximately 45 genera and 1,500 species, which are widespread in the tropical and subtropical Asia, Indian Ocean and Pacific basin, and neotropic regions (Plunkett et al., 2004; Wen et al., 2008). The majority of the Araliaceae members are woody species that exhibit high adaptability to both the sunlight and shade environments (Li and Wen, 2016). Yet, all the Araliaceae species possess relatively conserved floral morphology, *i*.*e*., mostly 5-merous flowers with inferior ovaries (Philipson, 1970). High diversity of the leaf morphology is observed, which varies from simple and palmately compound to variously pinnately compound (Philipson, 1970; Wen, 2001; Yi et al., 2004). These attributes offer the Araliaceae family an ideal system to explore how the woody species respond to variable light conditions. In this study, we investigated the phenotypic and transcriptional responses of the woody species *Fatsia japonica* under variable light conditions. As a typical shade-tolerant species within the family Araliaceae, *F. japonica* is widely planted in sheltered and semi-shaded spots of the urban areas as an ornamental shrub (Araus et al., 1986). Here we collected inflorescence and leaf samples from five natural micro-habitats with distinct light intensity conditions. Phenotypic and transcriptomic comparisons of these *F. japonica* samples provide an opportunity to address how a shade-tolerant woody species responds to heterogeneous micro-habitats.

To further examine whether similar shade-tolerant phenotypes have evolved in phylogenetically distant Araliaceae species, we performed common garden experiments for *F. japonica* together with four additional woody species (*Metapanax delavayi, Schefflera arboricola, Schefflera delavayi* and *Tetrapanax papyriferum*) under contrasting highlight and lowlight conditions. The five woody species were selected according to their leaf morphologies (*i*.*e*., narrow and broad leaf) and ecological habitats (*i*.*e*., shade-grown and sun-grown), as both features are the major determinants in response to changing light conditions. In parallel, we also surveyed transcriptomic features of the five species to evaluate whether they share similar gene regulatory networks to respond to variable light conditions. Our study will provide evolutionary and ecological perspectives on how the Araliaceae species have evolved advantage phenotypes to adapt to distinct ecological habitats.

## MATERIALS AND METHODS

### Plant materials and treatment conditions

Leaf and inflorescence samples of the *F. japonica* were collected from five different natural micro-habitats on the campus of Fudan University in Shanghai (N 31°20’31’’, E121°30’9’’) during October 2020-March 2021. Inflorescence samples were collected from shade and sunlight micro-habitats of the same population with five biological replicates, respectively. Likewise, leaf samples of the two pairs of micro-habitats (sunlight vs. shade and sunfleck vs. deepshade) were collected from the same individual with five biological replicates. In contrast, the leaf samples that grown under highlight condition were collected from an independent population in open environment (Table S1). To further examine whether the phenotypic responses are common in the family Araliaceae, we performed common garden experiment for one sun-grown (*T. papyrifer*) and four shade-grown (*F. japonica, M. delavayi, S*.*arboricola*, and *S. delavayi*) woody species under highlight (300 photosynthetically active radiation (PAR)) and lowlight (5 PAR) conditions, respectively (Table S1). All the common garden experiments were conducted in greenhouse (16-hour light at 22 °C /8-hour dark at 18°C) on the campus of Fudan University in Shanghai. Leaf samples were collected from the plants that grown under the two treatment conditions for 90 days (Table S1). Light quantity and quality were measured with the Plant Lighting Analyzer (PLA-30 V2.00, Hangzhou, China) (Table S1).

### Assessment of the plant performance under distinct light conditions

Light response curve was evaluated for the five woody Araliaceae species under the same condition in greenhouse using Li 6800 (Li-COR, Lincoln, 180 USA), with the sun-grown Arabidopsis (Columbia ecotype) and shade-grown herbal species *Panax ginseng* (family Araliaceae) as controls. Chlorophyll a and b content of the leaf samples were estimated for the five natural micro-habitats and common garden conditions, respectively. Total chlorophyll content was extracted using acetone and alcohol in certain ratio as reported (Zhao et al., 2018) and calculated by SynergyTM2 multi-detection microplate reader (Biotek, VT). Maximum quantum yield of PSII (Fv/Fm) of the common garden samples were evaluated using the program ImagingWin on plant phenotype imaging platform (Entoscan, Taiwan, China). Leaf morphologies of the five species were photographed by a Nikon COOLSCOPE II (Nikon Corporation, Tokyo, Japan). In addition, we also examined leaf anatomical structure for both the natural micro-habitat and common garden samples. The paraffin sectioning samples were obtained from > 15 biological replicates for each light condition. Fresh leaf tissues were dipped in formalin-acetic acid-alcohol and embedded in paraffin wax and stained with safranin-O and fast green. Leaf anatomical structure was recorded with a Nikon Eclipse E200 optical microscope (Nikon Corporation, Tokyo, Japan).

### RNA extraction, sequencing and Gene expression and functional enrichment

Leaf and inflorescence samples used for phenotyping were also subjected to extract RNAs. Total RNAs were extracted from liquid nitrogen frozen samples using RNA extraction kits (TIANGEN, Beijing, China). Quality of the extracted RNAs were evaluated with Nanodrop 2000C spectrophotometry (Thermo scientific, Waltham, USA) and sequenced using Illumina Novaseq 6000 platform (Illumina, San Diego, USA). Reference transcripts of the five species were assembled using Trinity v2.14.0 (Grabherr et al., 2011) with default parameters. Clean short Illumina reads of these leaf and inflorescence samples were mapped onto the assembled references using HISAT2 v2.1.0 (Kim et al., 2019). Raw mapped read counts were calculated by prepDE.py of the program StringTie (Pertea et al., 2015). Deseq2 was employed to identify differentially expressed genes (DEGs), with the cutoff “two-fold change and p value < 0.05”. Overall expression pattern of the samples collected from distinct light conditions were assessed using Principal Component Methods (PCA) in the R package ggplot2. Homologous genes of the DEGs among the five species were identified using Orthfinder (Emms et al., 2019), with the default parameters. KEGG functional enrichment of the DEGs were annotated at KAAS (https://www.genome.jp/tools/kaas/) (Moriya et al., 2007). Venn diagrams were shown by via R package Venn at https://cran.r-project.org/src/contrib/Archive. Heatmap of the DEGs were visualized using R package pheatmap at https://CRAN.R-project.org/package=pheatmap.

## RESULTS

### Physiological traits associated with the shade-tolerance

To determine whether the shade-grown species have evolved advantage phenotypes, we measured the light response curve of all the five woody Araliaceae species, including *F. japonica, M. delavayi, S. arboricola, S. delavayi* and *T. papyrifer*um, together with the Arabidopsis and *Panax ginseng* (ginseng) as the controls. Our comparisons revealed that both the sun-grown herb species Arabidopsis (700-800 μmol·m^−2^·s^−1^) and woody species *T. papyrifer*um (700-800 μmol·m^−2^·s^−1^) possessed obviously higher light saturation point than the five shade-grown species (from 300 to 500 μmol·m^−2^·s^−1^) (Figure 1a; Table S2). In particular, all the five shade-grown Araliaceae species showed higher assimilation ability of the net photosynthetic rate and lower dark respiration rate compared to the two sun-grown species.

**FIGURE 1.**
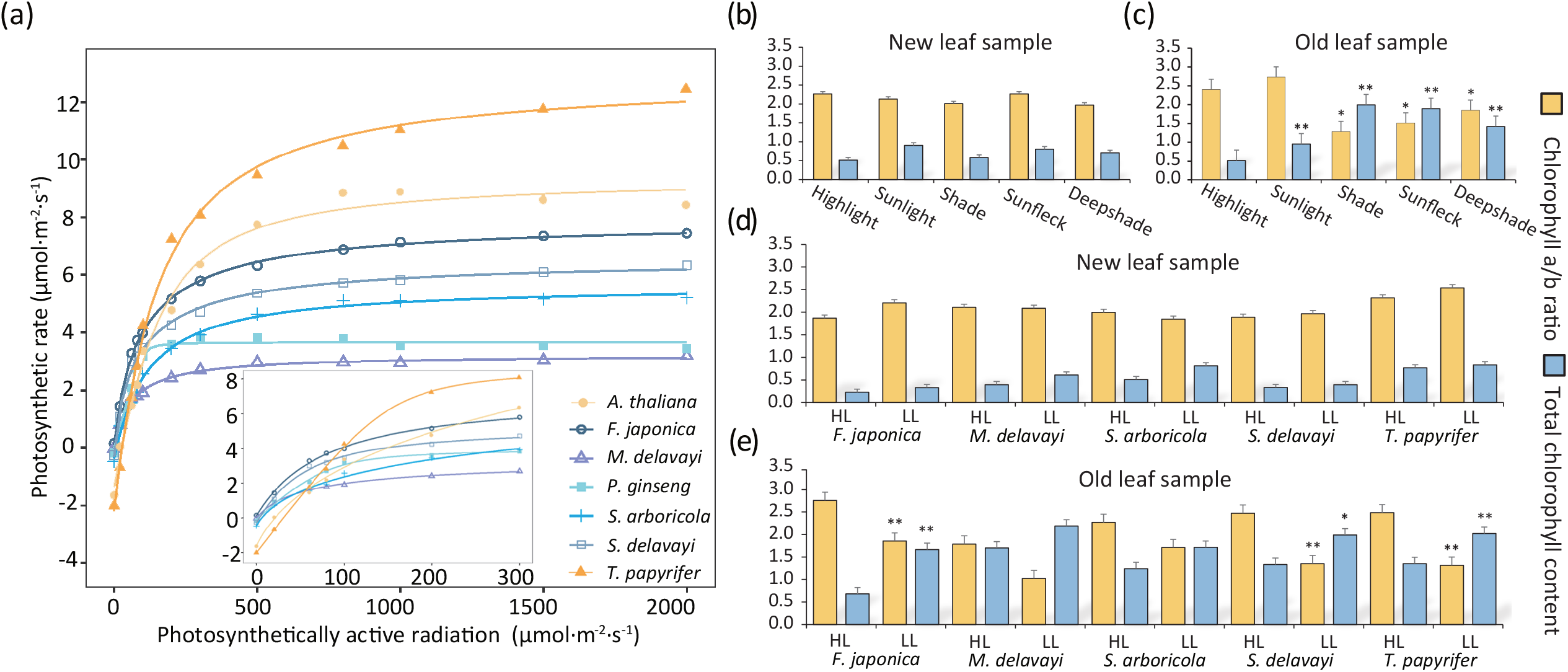
Light response curve and leaf chlorophyll content in the sun- and shade-grown species. (a) Colored lines represent the light response curve of the sun- and shade-grown species. X- and Y-axis are the photosynthetically active radiation (PAR) and photosynthetic rate, respectively. (b-c) Total leaf chlorophyll content (blue) and a/b ratio (orange) of the new and old leaf samples collected from the five micro-habitats of *Fatsia japonica*. (d-e) Total leaf chlorophyll content (blue) and a/b ratio (orange) of the new and old leaf samples collected from the common garden experiment of the five Araliaceae species. Significances of the total chlorophyll content and chlorophyll a/b ratio were performed among the leaf samples collected from the five micro-habitats of *Fatsia japonica*, as well as between the highlight and lowlight conditions of the five species. *, *Pr* value < 0.05, **, *Pr* value < 0.01.

The leaf chlorophyll content is a crucial physiological trait that determine photosynthesis efficiency of a plant under variable light conditions. We then measured total chlorophyll content and chlorophyll a/b ratio of the *F. japonica* leaf samples that grown naturally in five micro-habitats (highlight, sunlight, deepshade, shade and sunfleck). The new leaf samples collected from five micro-habitats showed no significant differences in total chlorophyll content and chlorophyll a/b ratio (bartlett-test, all *Pr* values > 0.05) (Figure 1b; Table S3). In contrast, the old leaf samples collected from the same individual of the five light conditions differed significantly in total chlorophyll content and chlorophyll a/b ratio (bartlett-test, all *Pr* values < 0.05) (Figure 1c; Table S3). In particular, the old leaf samples collected from the same individuals of the three shaded micro-habitats (deepshade, shade and sunfleck) accumulated more total chlorophyll content compared to the new leaf samples (Table S3). However, both the new and old leaf samples collected from the highlight and sunlight micro-habitats maintained similar level of total chlorophyll content (bartlett-test, all *Pr* values > 0.05).

In the common garden experiment, new leaf samples of the four shade-grown species showed similar chlorophyll a/b ratio (1.84-2.21) in both the highlight and lowlight conditions, but both are relatively lower than the new leaf samples of *T. papyrifer*um (sun-grown species) (2.32-2.54) (bartlett-test, all *Pr* values > 0.05) (Figure 1d; Table S3). In contrast, the old leaf samples collected from lowlight condition showed low chlorophyll a/b ratio compared to the highlight condition across all the five species (bartlett-test, three *Pr* values < 0.05) (Figure 1e; Table S3). For the total chlorophyll content, a general pattern was that both new and old leaf samples collected from the low light condition contained more total chlorophyll content compared to the high light condition (bartlett-test, three *Pr* values < 0.05) (Figure 1 d-e; Table S3). In particular, the decreased in chlorophyll a/b ratio in old leaf samples of the lowlight condition was mainly due to the accumulation of chlorophyll b content (Table S3). In addition, our estimations of the chlorophyll fluorescence parameter revealed that while the four shade-grown species possessed distinct leaf morphologies, they all maintained similar values of the maximum quantum yield of PS II (Fv/Fm) between the highlight and lowlight conditions (Figure 2; Table S4). In contrast, the sun-grown species *T. papyrifer*um exhibited significantly decreased in Fv/Fm value in lowlight condition. These attributes indicate that the four shade-grown species have evolved advantage phenotypes to cope with lowlight environments.

**FIGURE 2.**
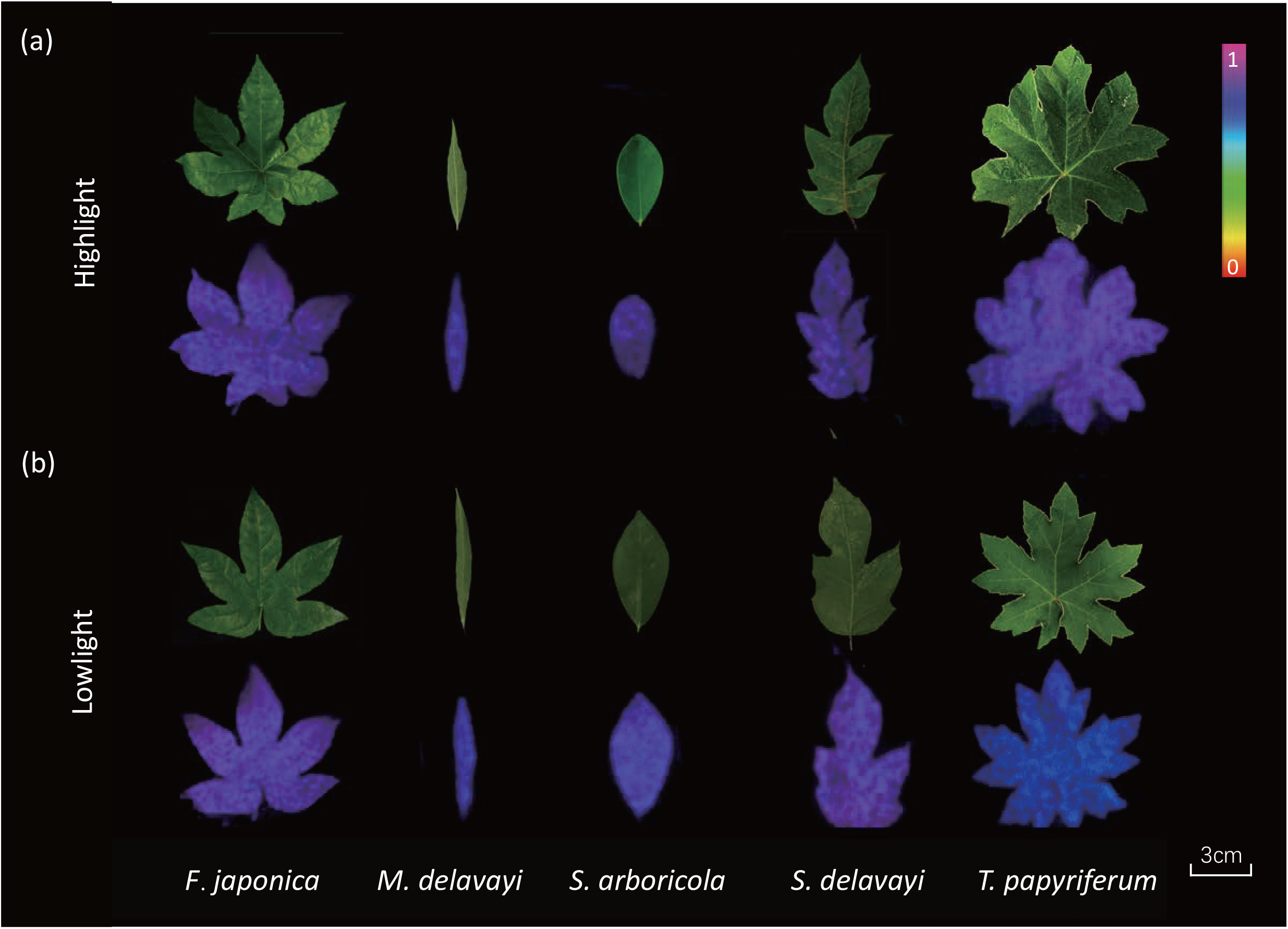
Chlorophyll fluorescence parameter of the five Araliaceae species grown under lowlight and highlight conditions. Leaf morphology and maximum quantum yield of PS II (Fv/Fm) of the five species under highlight (a) and lowlight (b) conditions, repsectively. All plants were grown under the two contrasting light conditions for three months. From left to right are *Fatsia japonica, Metapanax delavayi, Schefflera arboricola, Schefflera delavayi* and *Tetrapanax papyriferum*. The color bar on the upright corner represents the photosynthesis efficiency (Fv/Fm value = 0-1) from low (red) to high (purple).

### Phenotypic plasticity of the leaf anatomical structure

We examined leaf anatomical structures of the five Araliaceae species to assess whether the shade- and sun-grown woody plants showed different degrees of phenotypic plasticity. Among the *F. japonica* leaf samples, the palisade layer cells of the new leaf samples that grown under highlight and sunlight micro-habitats possessed closely packed cylindrical or rectangular cells (Figure 3a). In contrast, new leaf samples collected from the three shaded micro-habitats (deepshade, shade and sunfleck) have square and loosely packed palisade cells. In addition, the spongy layer cells of the three shade micro-habitats are in rounder shape with larger air pocket compared to the highlight and sunlight leaf samples (Figure 3a). The distinct leaf anatomical structures are more evident in the old leaf samples that collected from the five micro-habitats. In particular, both the new and old leaf samples grown under sunlight micro-habitat possessed multiple tightly arranged palisade layers.

**FIGURE 3.**
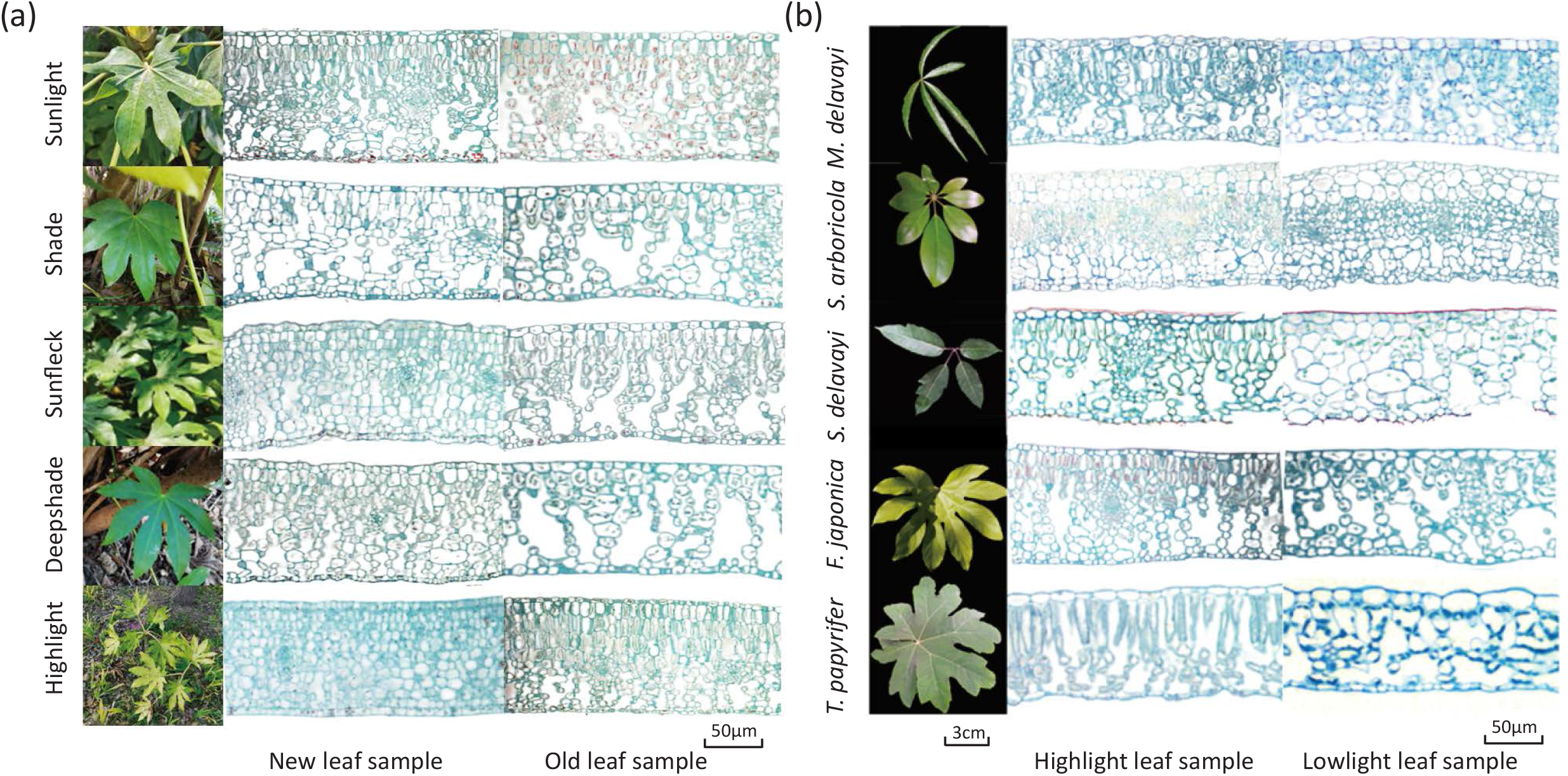
Leaf anatomical structure of the five species grown under distinct light conditions. (a) From top to bottom are the anatomical structure of the leaf samples collected from sunlight, shade, sunfleck, deepshade and highlight, respectively. From left to right are the five natural micro-habitats, anatomical structure of the new and old leaf samples, respectively. (b) From top to bottom are leaf samples of the five species (*Metapanax delavayi, Schefflera arboricola, Schefflera delavayi, Fatsia japonica* and *Tetrapanax papyriferum*). From left to right are the leaf morphology, anatomical structure of the new and old leaf samples collected from highlight and lowlight conditions, respectively. The bar on bottom is the exact size of the leaf sample.

We also compared the leaf anatomical structures of the common garden samples which were treated under the two contrasting light conditions for three months. Consistent with the above observations, both the new and old leaf samples that grown under highlight condition possessed cylindrical or rectangular palisade cells in all the four shade-grown species (Figure 3b). In contrast, palisade cells of the leaf samples collected from lowlight condition are round or square in shape (Figure 3b). It is notable that *T. papyrifer*um (sun-grown) showed a higher degree of developmental plasticity between the highlight and lowlight conditions compared to the four shade-grown species (Figure 3b).

### Transcriptional features under variable light conditions

The above phenotypic comparisons revealed that the four shade-grown species have evolved common shade-tolerance phenotypes (*i*.*e*., high photosynthesis efficiency) to cope with the shaded micro-habitats. We then asked whether these species also shared similar transcriptional features in response to variable light conditions. Of the *F. japonica* samples, our PCA analyses revealed that leaf samples collected from the five micro-habitats exhibited overall expression divergence, with the sunlight and highlight samples partially overlapped with each other but differing obviously from the other three shaded micro-habitats samples (deepshade, shade and sunfleck) (Figure 4a). Likewise, inflorescence samples collected from sunlight and shade micro-habitats not only exhibited distinct overall transcription patterns to each other but also differed obviously to all the leaf samples (Figure S1a). It is notable that although leaf samples of the two micro-habitat pairs (shade vs. sunlight and sunfleck vs. deepshade) were collected from the same individuals, they still possessed different degrees of gene expression divergence (Figure 4a). Likewise, the common garden samples that collected from highlight and lowlight conditions also showed overall gene expression divergence in all the five species (Figure S1b-f).

**FIGURE 4.**
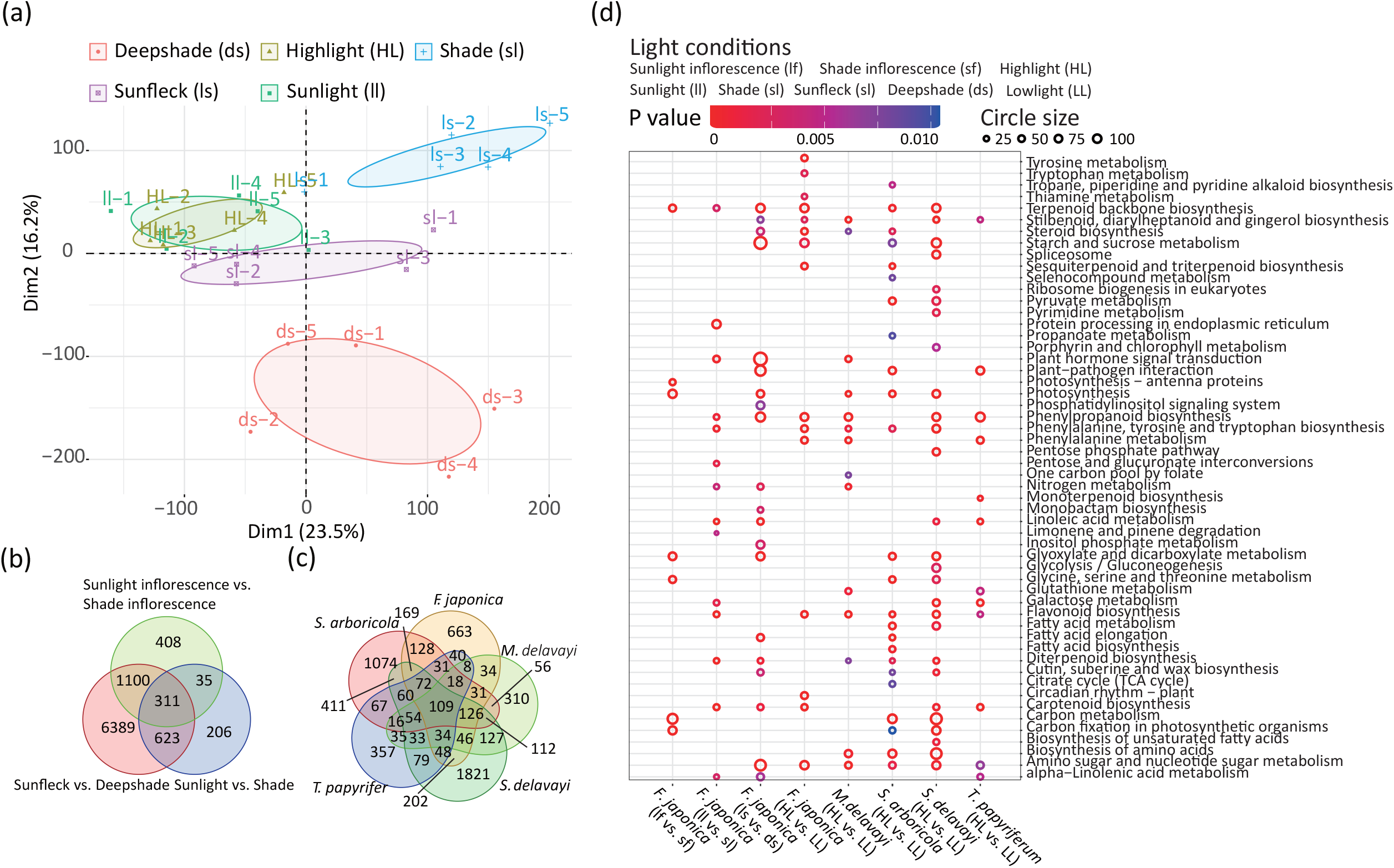
Overall transcriptomic features of the five species grown under distinct light conditions. (a) Expression pattern of the leaf samples collected from the five natural micro-habitats of *Fatsia japonica*. Samples collected from the same micro-habitat were marked with the same color. (b-c) Intersection analyses of the gene family groups that showed differential expression between the light conditions of the five species. The left Venn diagram is the common and unique gene family groups identified in leaf and inflorescence samples of the *Fatsia japonica*. The right diagram is the common and unique gene family groups identified between the highlight and lowlight conditions of the five species. Numbers in the Venn diagrams are the shared and unique gene family groups. (d) Functional enrichment of the DEGs identified in the five species. Each column represents the enrichment of the identified DEGs in each comparison. From left to right, the first three columns are the comparisons between different natural micro-habitats of the inflorescence and leaf samples of the *Fatsia japonica*. The last five columns are the comparisons between highlight (HL) and lowlight (LL) conditions of the common garden experiment of the five species. The colored bar on the top indicates significance of the functional enrichment. Names of each enriched term were shown on the right. Circle size indicate the number of gene in each enriched pathway.

We next identified genes that were differentially expressed in *F. japonica* among the five micro-habitats. Our analyses identified 1,463-14,902 (3.60%-35.14% of the total genes) DEGs corresponding to 1,061-8,423 family groups in the five species (Figure 4b and 4c; Table S6 and S7). Venn diagram revealed that a total of 11,864 family groups were involved in the response to distinct light conditions in the five species (Table S5). In the leaf and inflorescence samples of *F. japonica*, 311 (2.72%) of the 11452 family groups were commonly found in the comparisons of shade vs. sunlight and sunfleck vs. deepshade, respectively (Figure 4c). Likewise, 109 (1.71%) of 6371 family groups also shared in the common garden samples of the five species (Figure 4c).

Functional enrichment analyses of the KEGG showed that DEGs identified in leaf and inflorescence samples of *F. japonica* were functionally related to the pathways of plant photosynthesis (*i*.*e*., photosynthesis, carbon fixation and carotenoid biosynthesis), secondary metabolite (*i*.*e*., terpenoid backbone biosynthesis, diterpenoid and phenylpropanoid biosynthesis) and basic metabolisms (*i*.*e*., nitrogen, linoleic acid and glyoxylate and dicarboxylate metabolism) (Figure4d; Figure S2; Table S8). Likewise, DEGs characterized in the common garden samples were also functionally enriched in similar pathways, particularly those of related to photosynthesis (*i*.*e*., photosynthesis and carotenoid biosynthesis) and secondary metabolite (*i*.*e*., terpenoid backbone and phenylpropanoid biosynthesis). It is notable that the majority of the enriched KEGG pathways shared between the four shade-grown species and *T. papyrifer*um (sun-grown). Taken together, these findings suggest that similar gene regulatory networks might have played important roles in response to variable light conditions of the five species.

### Candidate genes involved in the responses to distinct light condition

Based on the analyses of gene functional enrichment detailed above, we further examined how the genes related to photosynthesis and photomorphogenesis expressed in the five species. Of the *F. japonica* samples, a general pattern is that the majority of these DEGs were differentially expressed in either different tissues (*i*.*e*., leaf vs. inflorescence) or micro-habitats (*i*.*e*., sunlight vs. shade light) (Figure 5a and b; Table S9-10). Among the photosynthesis-related genes, the majority of the cluster 1 genes (high expression in inflorescence tissue) were functionally related to carbon fixation and electron transport (Figure 5a; Table S9). Likewise, the cluster 2 and 3 genes (high expression in leaf tissue) were mainly involved in PS-I, PS-II, carbon fixation and electron transport. Notably, the cluster 3 genes (high expression in leaf tissue of the shade and deepshade samples) contained numerous antenna protein genes of the light-harvesting chlorophyll a/b-binding II (LHC-II) complex. In contrast, the cluster 2 genes (high expression in highlight and sunlight samples) were functionally correlated to non-photochemical quenching (NPQ) and antenna protein genes of the LCH-I complex.

**FIGURE 5.**
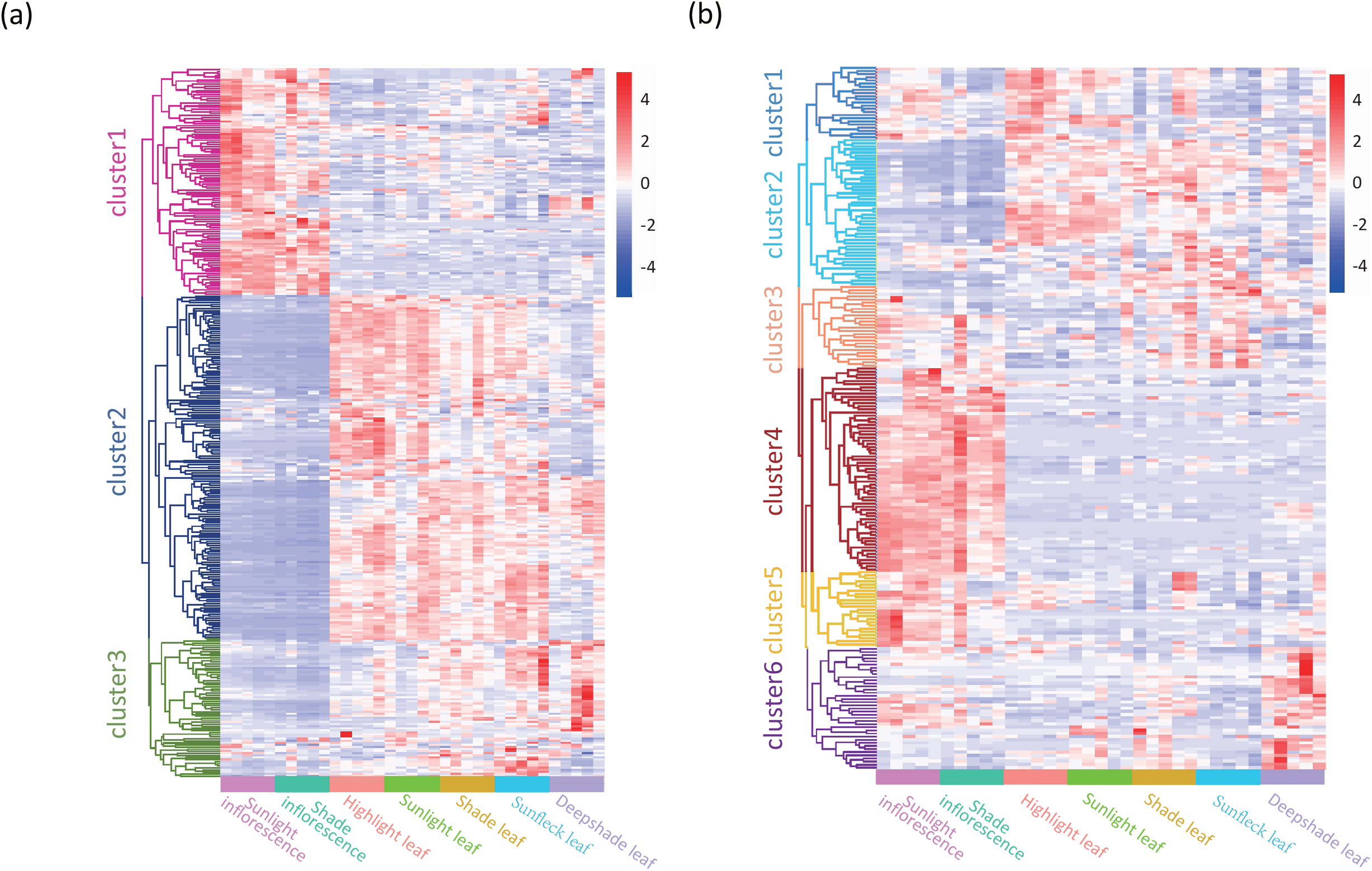
Transcriptional level of the photosynthesis- (a) and photomorphogenesis-related (b) genes in leaf and inflorescence samples collected from the five natural micro-habitats of *Fatsia japonica*. Each gene cluster is colored on the left of the heatmap. From left to right are the samples collected from sunlight inflorescence, shade inflorescence, highlight leaf, sunlight leaf, shade leaf, sunfleck leaf and deepshade leaf, respectively. Gene ID and exact expression values are shown in Table S9-10.

Similar patterns were also observed in the photomorphogenesis regulatory network genes where DEGs were identified in the comparisons of between the two tissues (leaf and inflorescence) and among the five micro-habitats (Figure 5b; Table S10). For example, high expression genes identified in inflorescence tissue (cluster 4 and cluster 5) were functionally related to the biosynthesis of phytohormones (*i*.*e*., auxin, BA and GA). In contrast, the cluster 2 genes (high expression in leaf tissue) included photoreceptors (*i*.*e*., *PHOT2* and *CRY2*) and regulatory genes (*i*.*e*., *PIF1* and *COP1*). In addition, our results also revealed that the phytochromes (*PHYB, PHYC* and *PHYE*) showed up-regulation in highlight samples in both inflorescence and leaf tissues compared to lowlight samples. In contrast, several genes related the biosynthesis of phytohormones (*i*.*e*., JA, BA and GA) and cell wall elongation were specifically up-regulated in deepshade sample of the leaf tissue.

Among the common garden samples, our analyses showed that some genes shared the same up- or down-regulation patterns in the highlight condition across the five species. For example, both the *PsbS* and *Psb28* that involved in the NPQ of chlorophyll fluorescence exhibited significant up-regulation in highlight condition in the leaf samples of the five species (Figure S3a; Table S11). Likewise, the photosynthetic electron transport gene *petJ* also showed significant up-regulation in highlight condition across the five species. In contrast, the majority of the LCH-I and LCH-II complex genes were down-regulated in highlight condition. In particular, several LHC-II complex genes (*i*.*e*., *LHCa5* and *LHCb1*) were specifically down-regulated in the sun-grown species *T. papyrifer*um. Among the photoreceptors, our results showed that several red (*i*.*e*., *PHYB, PHYC* and *PHYE*) and blue light (*i*.*e*., *CRY3* and *PHOT2*) receptors exhibited relatively higher level of up-regulation in the four shade-grown species compared to the sun-grown species (Figure S3b; Table S12). These observations suggest that these candidate genes may have been involved in the molecular responses of the five species under distinct light conditions.

## DISCUSSION

### Plant performance of the woody sun- and shade-grown species

Light gradients are ubiquitous in nature that affect plant anatomy, morphology, physiology and life history (Poorter et al., 2019). The sun-grown plants (*i*.*e*., Arabidopsis) adopt short-time responses, collectively termed shade-avoidance syndrome, to escape from shaded environments (Molina-Contreras et al., 2019). In contrast, shade-grown plants have evolved tolerant phenotype as a long-term strategy to survive in shaded habitats (Gommers et al., 2013). The carbon gain hypothesis proposes that shade-tolerant species have evolved advantage phenotypes (*i*.*e*., increased total chlorophyll content and high net photosynthetic rate) to maximize the photosynthesis efficiency and minimize respiration costs for maintenance (Givnish, 1988; Valladares & Niinemets, 2008; Kaiser et al., 2019). Indeed, it has been documented that, compared to Arabidopsis, the shade-grown herbal species *P. ginseng* (family Araliaceae) maintains high photosynthesis efficiency under shade conditions (Zhang et al., 2022). In this study, we compared the plant performance of one sun-grown and four shade-grown woody species within the family Araliaceae. Our phenotypic assessments demonstrated that the woody species inhabiting shade environments have also evolved fitness phenotypes (*i*.*e*., high maximum quantum yield of PS II, low light saturation point and dark respiration rate) to cope with lowlight habitats. Consistent with Arabidopsis, the sun-grown woody species (*T. papyrifer*um) exhibited significantly decreased photosynthesis efficiency under lowlight condition. These phenotypic features support the carbon gain hypothesis that both herbal and woody Araliaceae species have evolved high photosynthesis capacity during the adaptation to shaded habitats.

Chlorophyll a and b are two major pigments in the antenna complex that absorb sunlight and convert into chemical energy through photosynthesis (Jansson, 1994; Jansson, 1999). Here we showed that new leaf samples of *F. japonica* collecting from distinct micro-habitats possessed similar total chlorophyll content and chlorophyll a/b ratios. However, old leaf samples that grown under the shade micro-habitats (shade, deepshade and sunflect) accumulated more total chlorophyll content relative to both the highlight and sunlight conditions. In particular, the majority of increased total chlorophyll content is due to the accumulation of chlorophyll b content. Similar phenomenon was also observed in the common garden samples where leaf samples collected from lowlight condition accumulated more chlorophyll b in both the sun- and shade-grown species. Chlorophyll a is the principal pigment of photosynthesis, whereas chlorophyll b is an accessory pigment collecting energy in order to pass into chlorophyll a (Aronoff, 1950). The shade adapted chloroplast contain more chlorophyll b in order to increase the range of absorbed light wave length (Gommers et al., 2013). Yet, we did not find strong evidence that the increase in chlorophyll b is a specific adaptive strategy of the shade-grown species. It is clear that the accumulation of chlorophyll b is associated with the response to low light conditions across the five Araliaceae species.

The phenotypic plasticity hypothesis proposes that shade-tolerant species tend to show low degree of physiological and morphological plasticity under variable light conditions (Valladares et al., 2002). Indeed, phenotypic comparisons of the plant performance revealed that P. ginseng (shade-grown) exhibited apparently lower degree of phenotypic plasticity compared to Arabidopsis (sun-grown), such as plant morphology and flowering time (Zhang et al., 2022). Likewise, meta-analyses of numerous phenotypic traits also confirmed that woody and shade-grown herbal species are expected to exhibit low degree of phenotypic plasticity under changing light conditions, such as photosynthetic capacity and realized growth (Poorter et al., 2019). Here we showed that the five woody Araliaceae species are highly diversity in leaf morphology, such as simple leaf, palmately compound and pinnately compound leaf. Nevertheless, they all maintain highly similar leaf morphologies among distinct light conditions. It is notable that all the five species showed distinct leaf anatomical structures in the five micro-habitats and common garden conditions. In particular, the sun-grown species (*T. papyrifer*um) showed dramatic changes in leaf anatomical structure between highlight and lowlight conditions. Together, our results support that both the herbal and woody shade-grown species exhibited lower degree of phenotypic plasticity under variable light conditions compared to the sun-grown species.

### Transcriptional features of the sun- and shade-grown species

Molecular regulatory network of sun-grown plants (*i*.*e*., Arabidopsis) in response to changing light conditions have been well-documented (Casal, 2013). Several hypotheses related to the suppression of the SAS have been proposed to explain the molecular bases underpinning the shade-tolerant phenotype (Gommers et al., 2013; Leivar et al., 2012). In *Cardamine hirsuta*, up-regulation of the *PHYA* gene acts as an antagonistic factor to inhibit hypocotyl elongation (Molina-Contreras et al., 2019). Likewise, loss function of phytochrome-interacting factors (*PIFs*) together with differential regulation of photosynthesis-related genes are supposed to contribute to the adaptation of P. ginseng to shaded habitat (Zhang et al., 2022).

In this study, we aimed to address the transcriptional feature of woody species under variable light environments. Our analyses revealed that all the five Araliaceae species showed different overall gene expression patterns under distinct light conditions, even these samples collected from the same individuals (*i*.*e*., shade vs. sunlight and sunflect vs. deepshade). In particular, a large number of the DEGs that identified between distinct micro-habitats or light conditions shared by the five species. It suggests that both the sun- and shade-grown species may have adopted similar gene regulatory networks to respond to distinct light conditions. It is notable that these DEGs are functionally related to photosynthesis, carbon fixation and secondary metabolites. Given that phenotypic comparisons identified fitness traits in the shade-grown species (*i*.*e*., high photosynthesis efficiency), the identified DEGs in the five Araliaceae species are potentially associated with the phenotypic responses to different light conditions. For example, NPQ is a photoprotection mechanism that protects the photosynthetic machinery from damage under highlight intensities (Jahns & Holzwarth, 2012; Ruban, 2016). Our transcriptomic analyses of the *F. japonica* samples confirmed that genes involved in NPQ (*i*.*e*., *PsbS* and *Psb28*) are up-regulated in both the inflorescent and leaf tissues under highlight conditions. Likewise, these NPQ-related genes also showed high transcription level under highlight condition of the common garden samples in both the shade-grown and sun-grown species. It confirms that NPQ is a common photoprotection mechanism in both the shade- and sun-grown woody species. In addition, we also observed up-regulation of the LHC-I and II genes under low light conditions in the five species. The two light-harvesting systems consist of proteins (*i*.*e*., antenna proteins) and photosynthetic pigments (*i*.*e*., chlorophyll a and b) that absorb solar energy and transfer to photosynthetic reaction center (Boekema, 1995; Hankamer & Barber, 1997; Dekker & Boekema, 2005; Tikhonov, 2014). High transcriptional level of the LHC-I and II genes, together with the accumulated chlorophyll a and b, may increase the capacity for harvesting light in low light environments.

In Arabidopsis, photomorphogenesis regulatory network plays essential roles in determining plant performance under shade light conditions, such as elongation of elongation petiole and accelerated flowering (Pierik & Testerink, 2014; Fraser, et al, 2016). Here our results also revealed that transcriptional levels of the photomorphogenesis regulatory genes differed both between the leaf and inflorescence tissues and among the five micro-habitats. For example, both the red (*i*.*e*., *PHYB*) and blue light (*i*.*e*., *CRY1/2*) receptors are supposed to be up-regulated in Arabidopsis under shade light condition (Pedmale, et al, 2016). However, we only observed up-regulation of the red-light receptors (*i*.*e*., *PHYB, PHYC* and *PHYE*) in both the inflorescence and leaf tissues under highlight condition. As the PHYB is also a temperature sensor (Legris, et al, 2016; Qiu et al, 2019), it is likely that these red-light receptors are involved in the thermomorphogenesis under the highlight environments. Likewise, the majority of the blue-light receptors (*i*.*e*., *CRY1/2*) are not up-regulated in the shade light condition of the inflorescence and leaf tissues. Instead, these cryptochromes and phototropins (*i*.*e*., *PHOT1*) are highly transcribed in leaf tissue, especially for the highlight samples. Cryptochromes are blue-light receptors controlling a range of responses that serve to optimize the photosynthetic efficiency of plants, such as regulation of stomatal opening, chloroplast relocation movements, and phototropism (Christie & Murphy, 2013; Briggs, 2014; Kong & Wada, 2014; Inoue & Kinoshita, 2017). High expression level of these blue-light receptors in leaf tissue indicate that these genes may play important roles in regulating leaf development and photosynthesis efficiency. It is notable that the four shade-grown species possessed a up-regulation at the red- (*i*.*e*., *PHYB, PHYC* and *PHYE*) and blue-light (*i*.*e*., *CRY3* and *PHOT1*) receptors compared to the sun-grown species. It indicates the possibility that shade-grown species may have evolved a rapid or sensitive strategy in response to changing light environments. Unfortunately, as all the five woody Ararliaceae species are non-model species without reference genome, it limits the investigations of molecular regulation mechanism underlying this ecological adaptation. However, our study still provides evidence of the transcriptomic features of the shade- and sun-grown species under variable light environments.

## AUTHORS CONTRIBUTIONS

L.F.L. conceived this project and coordinated research activities; Y.Q.N. collected and maintained the plant materials; Y.Q.N., Y.X.Z. and X.F.W. conducted the data analyses; Y.Q.N., Y.X.Z., X.F.W., J.Y., Z.P.S., Y.G.W., W.J.Z., Z.H.W. and L.F.L. interpreted the data and participated in discussion; Y.Q.N., J.Y., Z.P.S., Y.G.W., W.J.Z, Z.H.W. and L.F.L. wrote the manuscript. All authors discussed the results and approved the manuscript.

## ACKNOWLEDGEMENTS

This work was financially supported by National Natural Science Foundation of China (31970235), China Postdoctoral Science Foundation Grant (2018M630400), and Shanghai Pujiang Program (19PJ1401500).

## CONFLICT OF INTEREST

Funding bodies had no role in the design of the study and in the collection, analysis and interpretation of data, or writing the manuscript. The authors declare that they have no competing interests.

## DATA AVAILABILITY STATEMENT

All data generated in this study were submitted to GenBank under the Bioproject numbers PRJNA845005.

## LEGEND

**FIGURE S1** Principal component analysis of the overall expression pattern of the five Araliaceae species. (a) Overall expression pattern of the leaf and inflorescence samples collected from the five natural micro-habitats of *Fatsia japonica*. (b-f) Overall expression pattern of the leaf samples collected from highlight and lowlight conditions of the common garden experiment of the five species.

**FIGURE S2** Functional enrichment of the differential expression gene identified in the five species. Each column represents the enrichment of the identified differential expression genes.

**FIGURE S3** Transcriptional level of the photosynthesis- (a) and photomorphogenesis-related (b) genes in leaf samples collected from the common garden experiment of the five species.

## Notes

### Competing Interest Statement

The authors have declared no competing interest.

## REFERENCES

Araus, J. L., Alegre, L., Tapia, L., Calafell, R., & Serret, M. D. (1986). Relationships between photosynthetic capacity and leaf structure in several shade plants. American Journal of Botany, 73(12), 1760-1770. https://doi.org/10.1002/j.1537-2197.1986.tb09708.x

Aronoff, S. (1950). Chlorophyll. The Botanical Review volume, 16, 525–588.

Augspurger, C. K. (1984). Light requirements of neotropical tree seedlings: a comparative study of growth and survival. Journal of Ecology, 72(3), 777–795. https://doi.org/10.2307/2259531

Ballare, C. L., & Austin, A. T. (2019). Recalculating growth and defense strategies under competition: key roles of photoreceptors and jasmonates. Narnia, 70(13), 3425–3434. https://doi.org/10.1093/jxb/erz237

Baltzer, J. L., & Thomas, S. C. (2007). Determinants of whole-plant light requirements in Bornean rain forest tree saplings. Journal of Ecology, 95(6), 1208–1221. https://doi.org/10.1111/j.1365-2745.2007.01286.x

Boekema E. J., Hankamer B., & Rögner M. (1995). Supramolecular structure of the photosystem II complex from green plants and cyanobacteria. Proceedings of the National Academy of Sciences, 92(1), 175–179. https://doi.org/10.1073/pnas.92.1.175

Bou-Torrent, J., Salla-Martret, M., Brandt, R., Musielak, T., Palauqui, J. C., Martínez-García, J. F., & Wenkei, S. (2012). ATHB4 and HAT3, two class II HD-ZIP transcription factors, control leaf development in Arabidopsis. Plant Signaling & Behavior, 7(11), 1382–1387. https://doi.org/10.4161/psb.21824

Canham, C. D., Kobe, R. K., Latty, E. F., & Chazdon, R. L. (1999). Interspecific and intraspecific variation in tree seedling survival: effects of allocation to roots versus carbohydrate reserves. Oecologia, 121(1), 1–11. https://doi.org/10.1007/s004420050900

Casal, J. J. (2013). Photoreceptor signaling networks in plant responses to shade. Annual Review of Plant Biology, 64(1), 403–427. http://doi.org/10.1146/annurev-arplant-050312-120221

Dekker J. P., & Boekema E. J. (2005). Supramolecular organization of thylakoid membrane proteins in green plants. Biochimica et Biophysica Acta, 1706(1-2), 12–39. https://doi.org/10.1016/j.bbabio.2004.09.009

Delagrange, S., Messier, C., Lechowicz, M. J., & Dizengremel, P. (2004). Physiological, morphological and allocational plasticity in understory deciduous trees: importance of plant size and light availability. Tree Physiology, 24(7), 775–784. https://doi.org/10.1093/treephys/24.7.775

Emms, D. M., & Kelly, S. (2019). OrthoFinder: phylogenetic orthology inference for comparative genomics. Genome Biology, 20, 238. https://doi.org/10.1186/s13059-019-1832-y

Fiorucci, A. S., & Fankhauser, C. (2017). Plant strategies for enhancing access to sunlight. Current Biology, 27(17), R931–R940. https://doi.org/10.1016/j.cub.2017.05.085

Fraser, D. P., Hayes, S., & Franklin, K.A. (2016). Photoreceptor crosstalk in shade avoidance. Current Opinion in Plant Biology, 33, 1–7. https://doi.org/10.1016/j.pbi.2016.03.008

Givnish, T. J. (1988). Adaptation to sun and shade: a whole-plant perspective. Australian Journal of Plant Physiology, 15(2), 63–92. http://doi.org/10.1071/pp9880063

Gommers, C. M., Keuskamp, D. H., Buti, S., Veen, H. V., Koevoets, I. T., Reinen, E., Voesenek, L., & Pierik, R. (2017). Molecular profiles of contrasting shade response strategies in wild plants: differential control of immunity and shoot elongation. The Plant Cell, 29(2), 331–344. http://doi.org/10.1105/tpc.16.00790

Gommers, C. M., Visser, E. J., St Onge, K. R., Voesenek, L. A., & Pierik, R. (2013). Shade tolerance: when growing tall is not an option. Trends in Plant Science, 18(2), 65–71. http://doi.org/10.1016/j.tplants.2012.09.008

Grabherr, M. G., Haas, B. J., Moran, Y., Levin, J. Z., Thompson, D. A.,Amit, I., Adiconis, X., Fan, L., Raychowdhury, R., Zeng, Q., Chen, Z., Mauceli, E., Hacohen, N., Gnirke, A., Rhind, N., Palma, F. D., Birren, B. W., Nusbaum, C., Lindblad-Toh, K., Friedman, N., & Regev, A. (2011). Full-length transcriptome assembly from RNA-Seq data without a reference genome. Nature Biotechnology, 29(7), 644–652. https://doi.org/10.1038/nbt.1883

Hankamer B., & Barber J. (1997). Structure and membrane organization of photosystem II in green plants. Annual Review of Plant Physiology and Plant Molecular Biology, 48(1), 641–671. https://doi.org/10.1146/annurev.arplant.48.1.641

Huber, M., Nieuwendijk, N. M., Pantazopoulou, C. K. & Pierik, R. (2021). Light signalling shapes plant-plant interactions in dense canopies. Plant, Cell & Environmont, 44, 1014-1029. https://doi.org/10.1111/pce.13912

Hudson, J. E., Levia, D. F., Hudson, S. A., Bais, H. P., & Legates, D. R. (2017). Phenoseasonal subcanopy light dynamics and the effects of light on the physiological ecology of a common understory shrub, Lindera benzoin. PLoS One, 12(10), e0185894. https://doi.org/10.1371/journal.pone.0185894

Jahns, P., & Holzwarth, A. R. (2012). The role of the xanthophyll cycle and of lutein in photoprotection of photosystem II. Biochimica et Biophysica Acta-Bioenergetics, 1817(1), 182–193. https://doi.org/10.1016/j.bbabio.2011.04.012

Jansson, S. (1994). The light-harvesting chlorophyll a/b-binding proteins. Biochimica et Biophysica Acta, 1184(1), 1–19. https://doi.org/10.1016/0005-2728(94)90148-1

Jansson, S. (1999). A guide to the Lhc genes and their relatives in Arabidopsis. Trends in Plant Science, 4(6), 236–240. https://doi.org/10.1016/S1360-1385(99)01419-3

Kaiser, E., Galvis, V. C., & Armbruster, U. (2019). Efficient photosynthesis in dynamic light environments: a chloroplast’s perspective. Biochemical Journal, 476(19), 2725–2741. http://doi.org/10.1042/BCJ20190134

Kim, D., Paggi, J. M., Park, C., Bennett, C., & Salzberg, S. L. (2019). Graph-based genome alignment and genotyping with HISAT2 and HISAT-genotype. Nature Biotechnology, 37(8), 907–915. http://doi.org/10.1038/s41587-019-0201-4

Kitajima, K. (1994). Relative importance of photosynthetic traits and allocation patterns as correlates of seedling shade tolerance of 13 tropical trees. Oecologia, 98(3-4), 419–28. http://doi.org/10.1007/BF00324232

Kobe, R. K. (1997). Carbohydrate allocation to storage as a basis of interspecific variation in sapling survivorship and growth. Oikos, 80, 226–233. https://doi.org/10.2307/3546590

Lau, O. S., & Deng, X. W. (2010). Plant hormone signaling lightens up: integrators of light and hormones. Current Opinion in Plant Biology, 13(5), 571–577. https://doi.org/10.1016/j.pbi.2010.07.001

Legris, M., Klose, C., Burgie, E. S., Rojas C. C. R., Neme, M., Hiltbrunner, A., Wigge, P. A., Schäfer, E., Vierstra, R. D. & Casal, J. J. (2016). Phytochrome B integrates light and temperature signals in Arabidopsis. Science, 354(6314), 897. https://www.science.org/doi/10.1126/science.aaf5656

Leivar, P., Monte, E., Cohn, M. M., & Quail, P. H. (2012). Phytochrome signaling in green Arabidopsis seedlings: impact assessment of a mutually negative phyB-PIF feedback loop. Molecular Plant, 5(3), 734–749. https://doi.org/10.1093/mp/sss031

Li, R., & Wen, J. (2016). Phylogeny and diversification of Chinese Araliaceae based on nuclear and plastid DNA sequence data. Journal of Systematics and Evolution, 54(4), 453–467. https://doi.org/10.1111/jse.12196

Liu, Y., Jafari, F. & Wang, H. (2021). Integration of light and hormone signaling pathways in the regulation of plant shade avoidance syndrome. aBIOTECH, 2(2), 131–14. https://doi.org/10.1007/s42994-021-00038-1

Molina-Contreras, M. J., Paulisic, S., Then, C., Moreno-Romero, J., Pastor-Andreu, P., Morelli, L., Roig-Villanova, I., Jenkins, H., Hallab, A., Gan, X., Gomez-Cadenas, A., Tsiantis, M., Rodríguez-Concepción, M., & Martinez-Garcia, J. (2019). Photoreceptor activity contributes to contrasting responses to shade in cardamine and Arabidopsis seedlings. The Plant Cell, 31(11), 2649–2663. https://doi.org/10.1105/tpc.19.00275

Moriya, Y., Itoh, M., Okuda, S., Yoshizawa, A C., & Kanehisa, M. (2007). KAAS: an automatic genome annotation and pathway reconstruction server. Nucleic Acids Research, 35, 182–185. https://doi.org/10.1093/nar/gkm321

Pedmale, U. V., Huang, S. S. C., Zander, M., Cole, B. J., Hetzel, J., Ljung, K., Reis, P., Sridevi, P., Nito, K., Nery, J. R., Ecker, J. R., & Chory, J. (2016). Cryptochromes interact directly with PIFs to control plant growth in limiting blue light. Cell, 164(1-2), 233-245. https://doi.org/10.1016/j.cell.2015.12.018

Pertea, M., Pertea, G. M., Antonescu, C. M., Chang, T. C., Mendell, J. T., & Salzberg, S. L. (2015). StringTie enables improved reconstruction of a transcriptome from RNA-seq reads. Nature Biotechnology, 33(3), 290–295. https://doi.org/10.1038/nbt.3122

Philipson, W. R. (1970). Constant and variable features of the Araliaceae. Botanical Journal of the Linnean Society, 63, 87–100.

Pierik, R., & Testerink, C. (2014). The art of being flexible: how to escape from shade, salt, and drought. Plant physiology, 166(1), 5–22. https://doi.org/10.1104/pp.114.239160

Plunkett G. M., Wen, J., & Lowry, P. P. (2004). Infrafamilial classifications and characters in Araliaceae: Insights from the phylogenetic analysis of nuclear (ITS) and plastid (trnL-trnF) sequence data. Plant Systematics and Evolution, 245(1-2), 1–39. https://doi.org/10.1007/s00606-003-0101-3

Poorter, H., Niinemets, U., Ntagkas, N., Siebenkas, A., Maenpaa, M., Matsubara, S. & Pons, T. (2019). A meta-analysis of plant responses to light intensity for 70 traits ranging from molecules to whole plant performance. New Phytologist, 223, 1073–1105. https://doi.org/10.1111/nph.15754

Qiu, Y. J., Li, M., Kim, R. J., Moore C. M., & Chen, M. (2019). Daytime temperature is sensed by phytochrome B in Arabidopsis through a transcriptional activator HEMERA. Nature Communications, 10(1), 140. https://doi.org/10.1038/s41467-018-08059-z

Ruban, A. V. (2016). Nonphotochemical chlorophyll fluorescence quenching: mechanism and effectiveness in protecting plants from photodamage. Plant Physiology, 170(4), 1903–1916. https://doi.org/10.1104/pp.15.01935

Sullivan, J. A., & Deng, X. W. (2003). From seed to seed: the role of photoreceptors in Arabidopsis development. Developmental Biology, 260(2), 289–297. http://doi.org/10.1016/s0012-1606(03)00212-4

Sultan, S. E. (2000). Phenotypic plasticity for plant development, function and life history. Trends in Plant Science, 5(12), 537–542. https://doi.org/10.1016/S1360-1385(00)01797-0

Valcarcel, V., & Wen, J. (2019). Chloroplast phylogenomic data support Eocene amphi-Pacific early radiation for the Asian Palmate core Araliaceae. Journal of Systematics and Evolution, 57(6), 547–560. https://doi.org/10.1111/jse.12522

Tikhonov, A. N. (2014). The cytochrome b6f complex at the crossroad of photosynthetic electron transport pathways. Plant Physiology & Biochemistry, 81, 163–183. https://doi.org/10.1016/j.plaphy.2013.12.011

Valladares, F., Chico, J. M., Aranda, I., Balaguer, L., Dizengremel, P., Manrique, E., & Dreyer, E. (2002). The greater seedling high-light tolerance of Quercus robur over Fagus sylvatica is linked to a greater physiological plasticity. Trees, 16(6), 395–403. https://doi.org/10.1007/s00468-002-0184-4

Valladares, F., & Niinemets, U. (2008). Shade tolerance, a key plant feature of complex nature and consequences. Annual Review of Ecology, Evolution, and Systematics, 39, 237–257. http://doi.org/10.1146/annurev.ecolsys.39.110707.173506

Wen, J. (2001). Evolution of the Aralia–Panax complex (Araliaceae) as inferred from nuclear ribosomal its sequences. Edinburgh Journal of Botany, 58(2), 243–257. http://doi.org/10.1017/S0960428601000610

Wen, J., Zhu, Y. P., Lee, C. H., Widjaja, E., & Saw, L. G. (2008). Evolutionary relationships of Araliaceae in the Malesian region: A preliminary analysis. Acta Botanica Yunnanica, 30(4), 391–399. https://doi.org/10.3724/SP.J.1143.2008.00391

Yi, T. S., Lowry, P. P., Plunkett, G. M., & Wen, J. (2004). Chromosomal evolution in Araliaceae and close relatives. Taxon, 53(4), 987–1005. https://doi.org/10.2307/4135565

Zhang, Y. X., Niu, Y. Q., Wang, X. F., Wang Z. H., Wang, M. L., Yang, J., Wang, Y. G., Zhang, W. J., Song, Z. P., & Li, L. F. (2022). Phenotypic and transcriptomic responses of the sun- and shade-loving plants to sunlight and dim-light conditions. https://doi.org/10.1101/2022.01.28.477942

