## Supplemental Figures for "Phenotypic and transcriptional features of the Araliaceae species under distinct light environments"

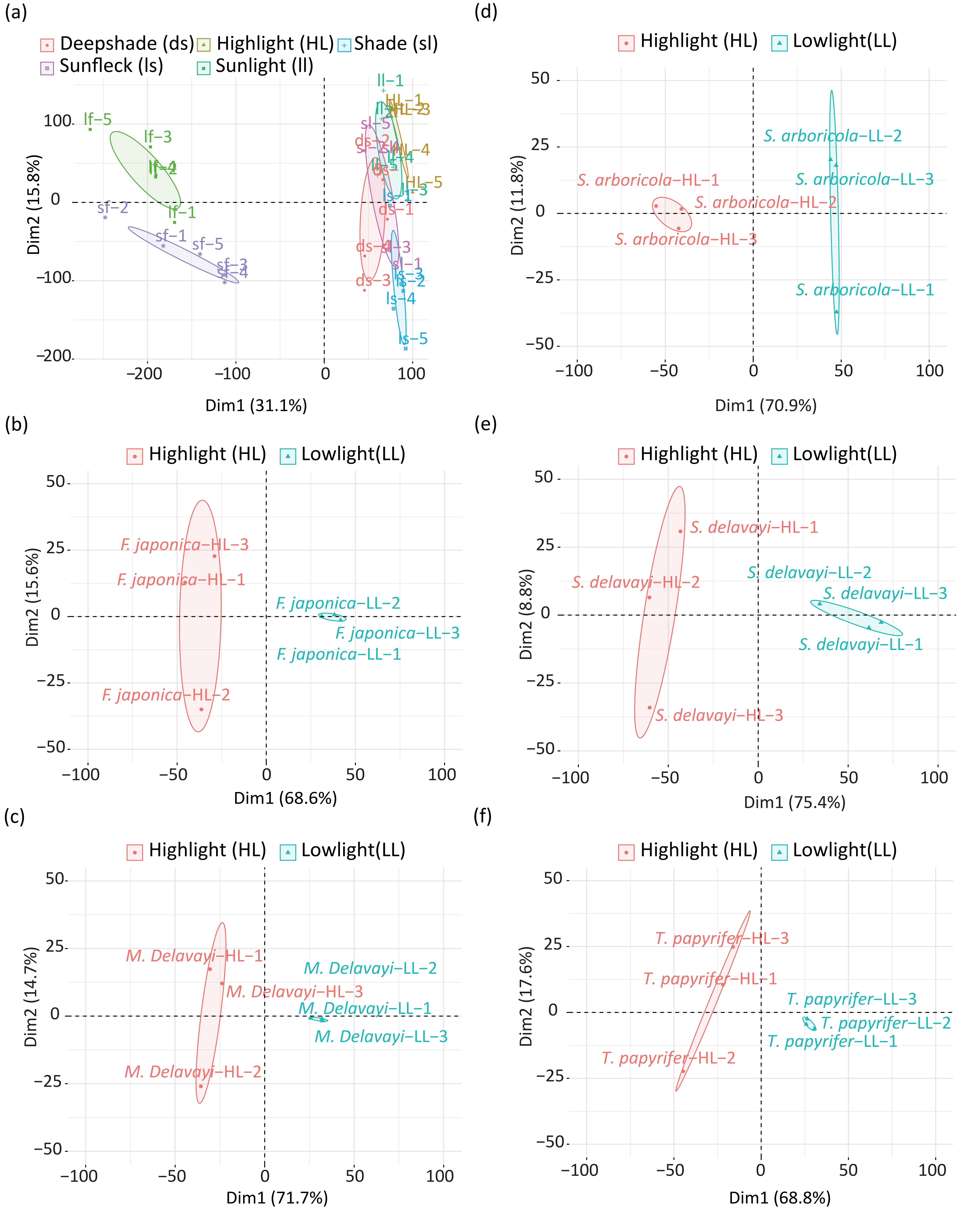


**FIGURE S1 Principal component analysis of the overall expression pattern of the five Araliaceae species.** (a) Overall expression pattern of the leaf and inflorescence samples collected from the five natural micro-habitats of *Fatsia japonica*. (b-f) Overall expression pattern of the leaf samples collected from highlight and lowlight conditions of the common garden experiment of the five species.

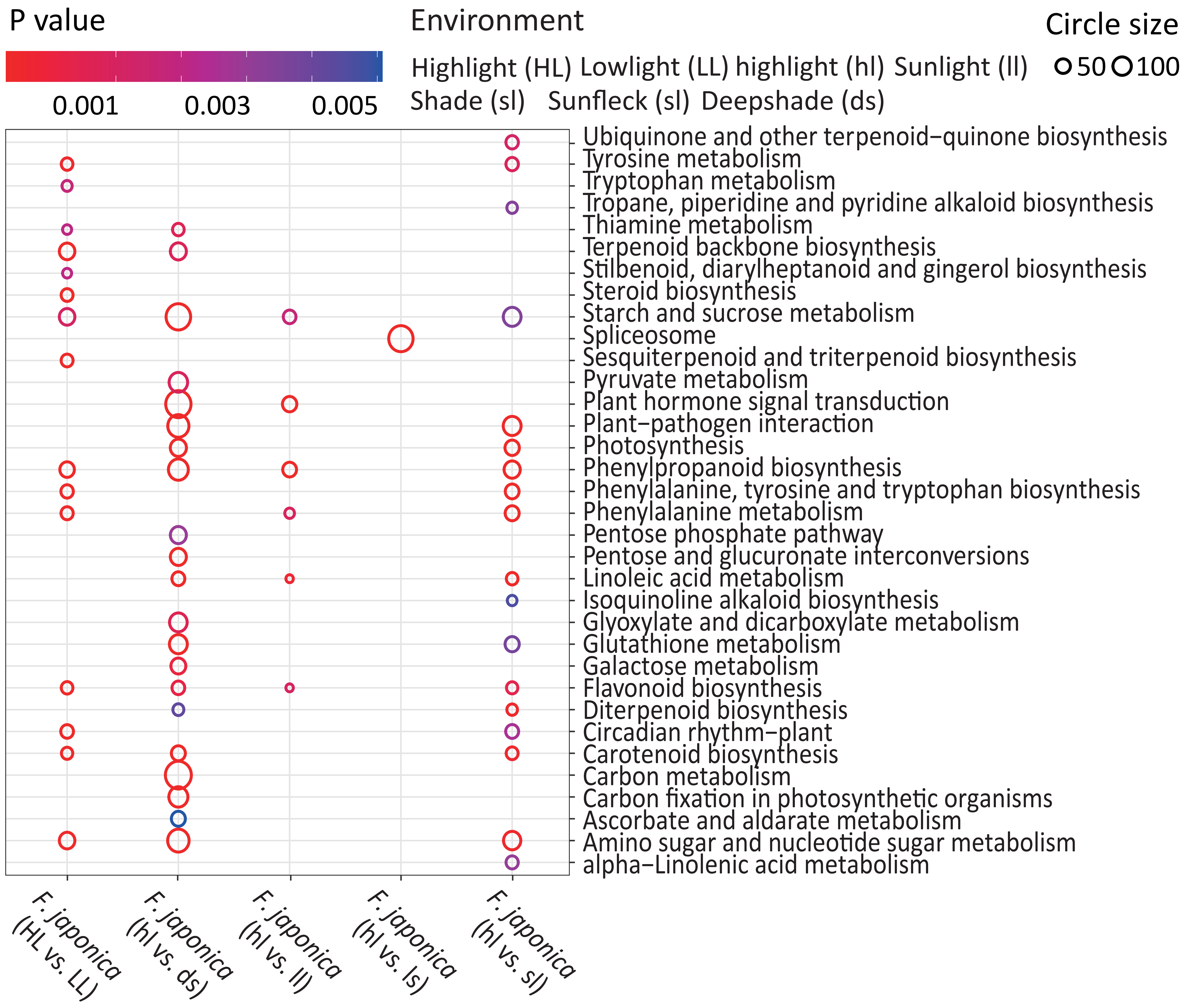


**FIGURE S2 Functional enrichment of the differential expression gene identified in the five species.** Each column represents the enrichment of the identified differential expression genes.

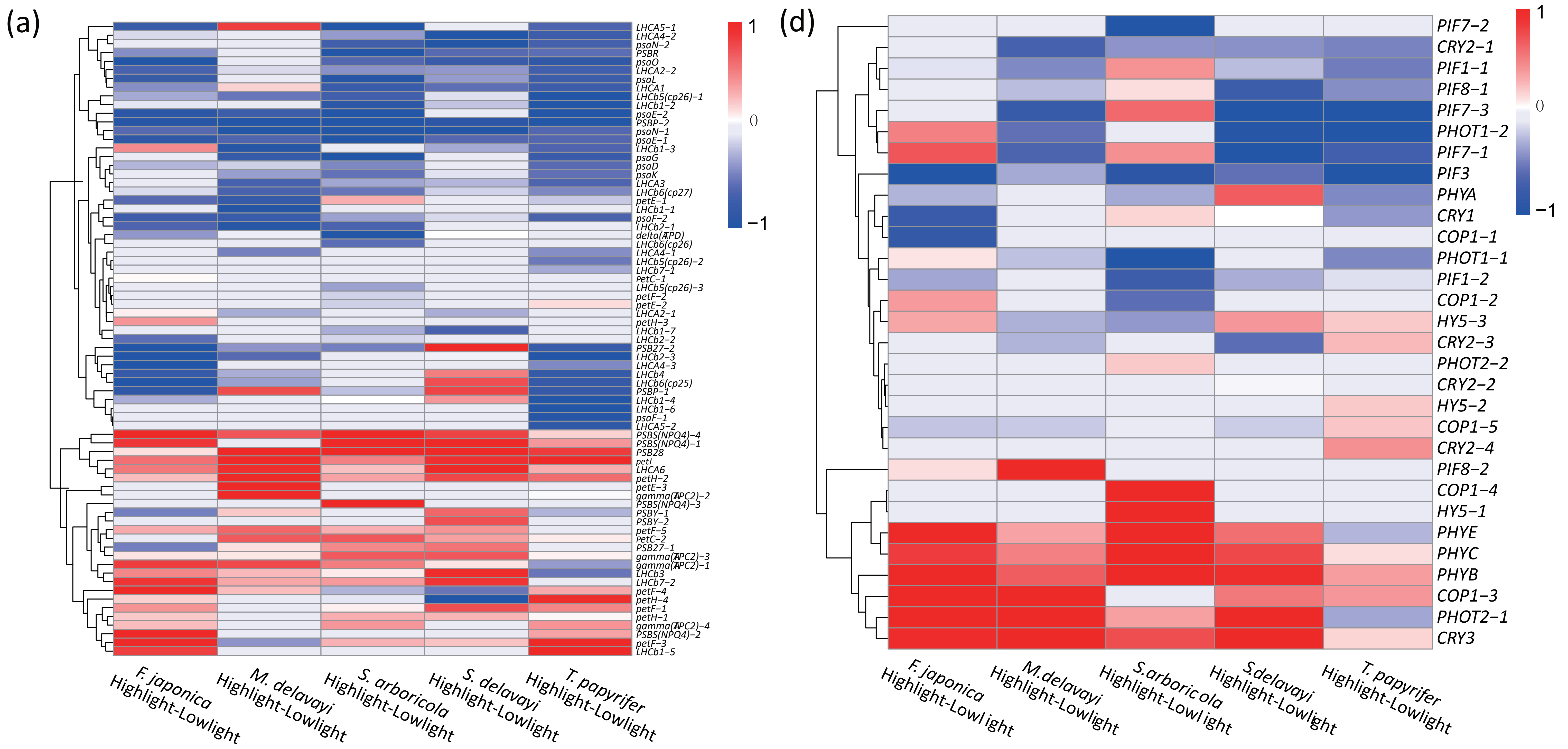


**FIGURE S3** Transcriptional level of the photosynthesis- (a) and photomorphogenesis-related (b) genes in leaf samples collected from the common garden experiment of the five species.
